# Multidimensional and Multiresolution Ensemble Networks for Brain Tumor Segmentation

**DOI:** 10.1101/760124

**Authors:** Gowtham Krishnan Murugesan, Sahil Nalawade, Chandan Ganesh, Ben Wagner, Fang F. Yu, Baowei Fei, Ananth J. Madhuranthakam, Joseph A. Maldjian

## Abstract

In this work, we developed multiple 2D and 3D segmentation models with multiresolution input to segment brain tumor components, and then ensembled them to obtain robust segmentation maps. This reduced overfitting and resulted in a more generalized model. Multiparametric MR images of 335 subjects from BRATS 2019 challenge were used for training the models. Further, we tested a classical machine learning algorithm (xgboost) with features extracted from the segmentation maps to classify subject survival range. Preliminary results on the BRATS 2019 validation dataset demonstrasted this method can achieve excellent performance with DICE scores of 0.898, 0.784, 0.779 for whole tumor, tumor core and enhancing tumor respectively and accuracy 34.5 % for survuval prediction.

## 1. Introduction

Brain Tumors account for 85% to 90% of all primary CNS tumors. The most common type of primary brain tumors are gliomas which can be further classified into High Grade Gliomas (HGG) and low grade Gliomas (LGG) based on their aggressiveness. Magnetic Resonance Imaging (MRI) is a widely used technique in diagnosis and clinical treatment of gliomas. Despite being a standard imaging modality for delineating tumors and treatment planning, using MRI for segmenting brain tumors remains a challenging task due to the high variation of brain tumor shape, size, location, and particularly due to the subtle intensity changes of tumor regions relative to the surrounding normal tissue. Consequently, manual tumor contouring is both time-consuming, and subject to large inter- and intra-observer variability. Semi- or fully-automated brain tumor segmentation methods could circumvent this variability for better patient management (Zhuge, Krauze et al. 2017). As a result, developing automated, semi-automated and interactive segmentation methods for brain tumor has huge clinical implications, however, this remains highly challenging. Efficient deep learning algorithms to segment brain tumors into their subcomponents help in early clinical diagnosis, treatment and follow-up of individual patients (Saouli, Akil et al. 2018).

The multimodal Brain Tumor segmentation Benchmark (BRATS) dataset provided a general platform by outsourcing a unique brain tumor dataset with known ground truth segmentations, done manually by experts (Menze, Jakab et al. 2014). Several advanced deep learning algorithms were developed on this unique platform provided by BRATS and benchmarked against common datasets allowing general comparisons between them. CNN-based methods have shown advantages with respect to learning the hierarchy of complex features and have performed best in the recent BRATS challenges. U-net(Ronneberger, Fischer et al. 2015) based network architectures have beenused for segmenting complex brain tumor structures. Pereira et al. developed a 2D CNN method with two CNN architectures for High Grade and Low Grade Gliomas separately and combined the outputs in the post processing steps (Pereira, Pinto et al. 2016). Havaei et al. developed a multiresolution cascaded CNN architecture with two pathways, each of which takes different 2D patch sizes with four MR sequences as channels (Havaei, Davy et al. 2017). The BRATS 2018 top performer developed a 3D decoder encoder style CNN architecture with inter-level skip connections to segment the tumor (Myronenko 2018). In addition to the decoder part, a Variation Autoencoder (VAE) was included to add reconstruction loss to the model.

In this study we propose to ensemble output from Multiresolution and Multidimensional models to obtain robust tumor segmentations. We utilized off the shelf model architectures (DensNET-169, SERESNEXT-101 and SENet-154) to perform segmentation using 2D inputs. We also implemented a two and three dimensional Residual Inception Densenet (RID) network to perform tumor segmentation with patch based inputs (64×64 and 64×64×64). The outputs from the model trained on different resolutions and dimensions were combined to eliminate false positives and post-processed using cluster analysis to obtain the final outputs.

## 2 Materials and Methods

### 2.1 Data and Preprocessing

The Brats 2019 dataset set included a total of 335 multi-institutional subjects(Menze, Jakab et al. 2014, Bakas, Akbari et al. 2017, Bakas, Akbari et al. 2017, Bakas, Akbari et al. 2017, Bakas, Akbari et al. 2017). It included 259 HGG subjects and 76 LGG subject. The standard preprocessing steps by the BRATS organizers on all MR images included co-registration to an anatomical template(Rohlfing, Zahr et al. 2010), resampling to isotropic resolution (1 mm3) and skull-stripping(Bakas, Reyes et al. 2018). Additional preprocessing steps included N4 bias field correction(Tustison, Cook et al. 2014) for removing the RF inhomogeneity and normalizing the multi parametric MR images to zero mean and unit variance.

The purpose of the survival prediction task is to predict the overall survival of the patient based on the multiparametric pre-operative MR imaging features in combination with the segmented tumor masks. Survival prediction based on only imaging based features (with age and resection status) is a difficult task based (. Additional information such as histopathological imaging, genomic information, radiotracer based imaging and other non imaging feature can be used to improve over all survival prediction. Pooya et.. al. (Mobadersany, Yousefi et al. 2018) reported better accuracy by combining genomic information and histopatholical images together to form a genomic survival convolutional neural network architecture (GSCNN model). Several studies have reported predicting overall survival for cerebral gliomas using ^11^C-acetate, and ^18^F-FDG in PET/CT scans (Tsuchida, Takeuchi et al. 2008, Yamamoto, Nishiyama et al. 2008, Kim, Kim et al. 2018).

### 2.2 Network Architecture

We trained several models to segment tumor components. All network architectures used for the segmentation task, except Residual Inception dense Network, were imported using Segmentation models, a python package(Yakubovskiy 2019). The models selected for the purpose of brain tumor segmentation had different backbones (DenseNet-169(Huang, Liu et al. 2017), SERESNEXT-101(Chen, Fan et al. 2018) and SENet-154(Hu, Shen et al. 2018)). The DenseNet architecture has shown some promising results in medical data classification and segmentation tasks (Islam and Zhang 2017, Chen, Wu et al. 2018, Dolz, Gopinath et al. 2018). The DenseNet model has advantages in feature propagation from one dense block to the next and overcomes the problem of the vanishing gradient (Huang, Liu et al. 2017). The squeeze and excitation block was designed to improve the feature propagation by enhancing the interdependencies between features for the classification task. This helps in propagating more useful features to the next block and suppressing less informative features. This network architecture was the top performer at the ILSVC 2017 classification challenge. SENet-154 and SE-ResNeXt-101 haves more parametersandis computationally expensive but has shown good results on the ImageNet classification tasks(Hu, Shen et al. 2018). Three of the proposed models were ensembled to obtain the final results. All of these models from the Segmentation Models package were trained with a 2D Axial Slice size of 240×240 (Fig.1).

**Figure 1:**
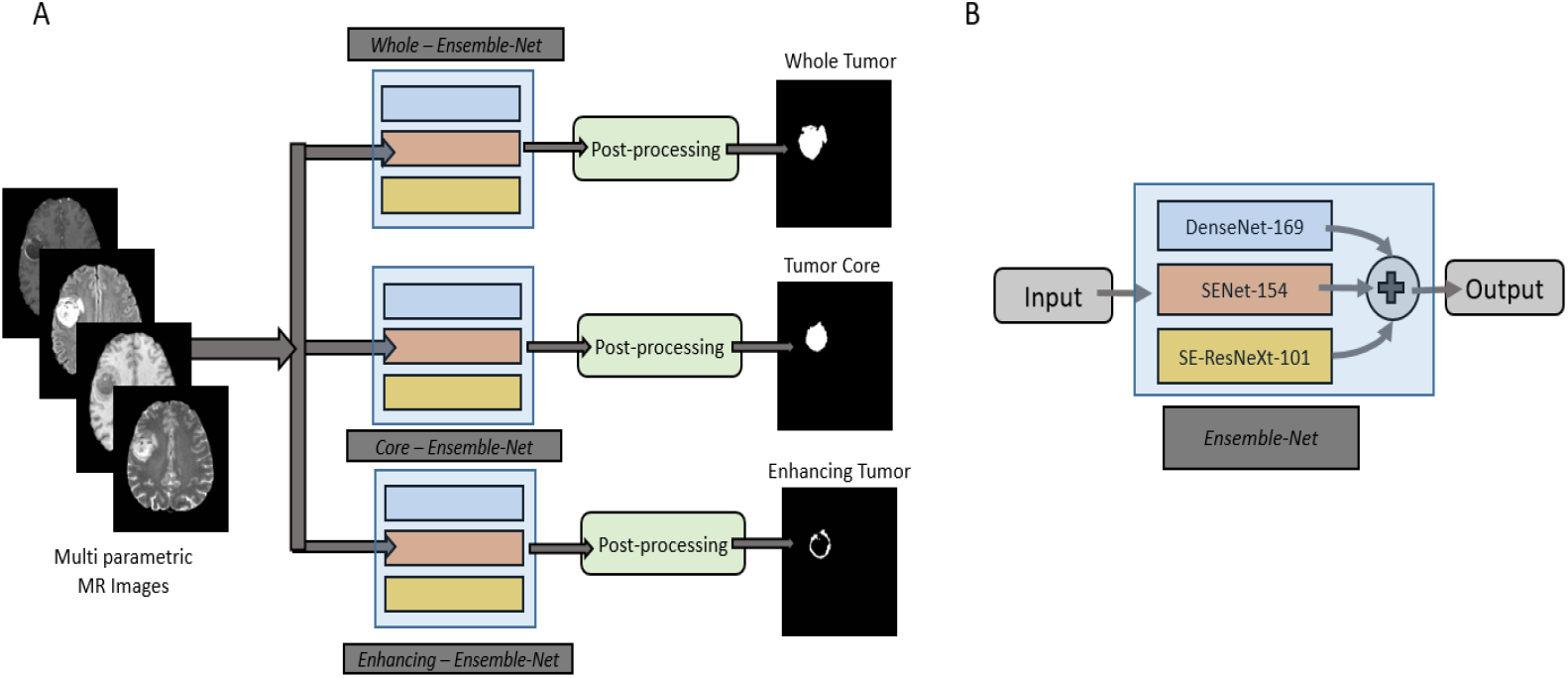
A. Ensemble of Segmentation models. (DenseNET-169, SERESNEXT-101 and SE-Net-154). B. Ensemble methodology used to combine the outputs from Segmentation Models to produce output segmentation maps

The Residual Inception Dense Network (RID) was first proposed and developed by Khened et al for cardiac segmentation. We incorporated our own implementation of the RID network in Keras with a Tensorflow backend (Figure.2). In the Densenet architecture, the GPU memory footprint increases with the number of feature maps of larger spatial resolution. The skip connections from the down-sampling path to the up-sampling path use element-wise addition in this model instead of the concatenation operation in Densenet, to mitigate feature map explosion in the up-sampling path. For the skip connections, a projection operation was done using BN-1 × 1-convolution-dropout to match the dimensions for element-wise addition (Figure.3). These additions to the Densenet architecture help in reducing the parameters and the GPU memory footprint without affecting the quality of segmentation output. In addition to performing dimension reduction, the projection operation helps in learning interactions of cross channel information (Lin, Chen et al. 2013) and faster convergence. Further, the initial layer of the RID networks includes parallel CNN branches similar to the inception module with multiple kernels of varying receptive fields, which help in capturing view-point dependent object variability and learning relations between image structures at multiplescales.

**Fig. 2.**
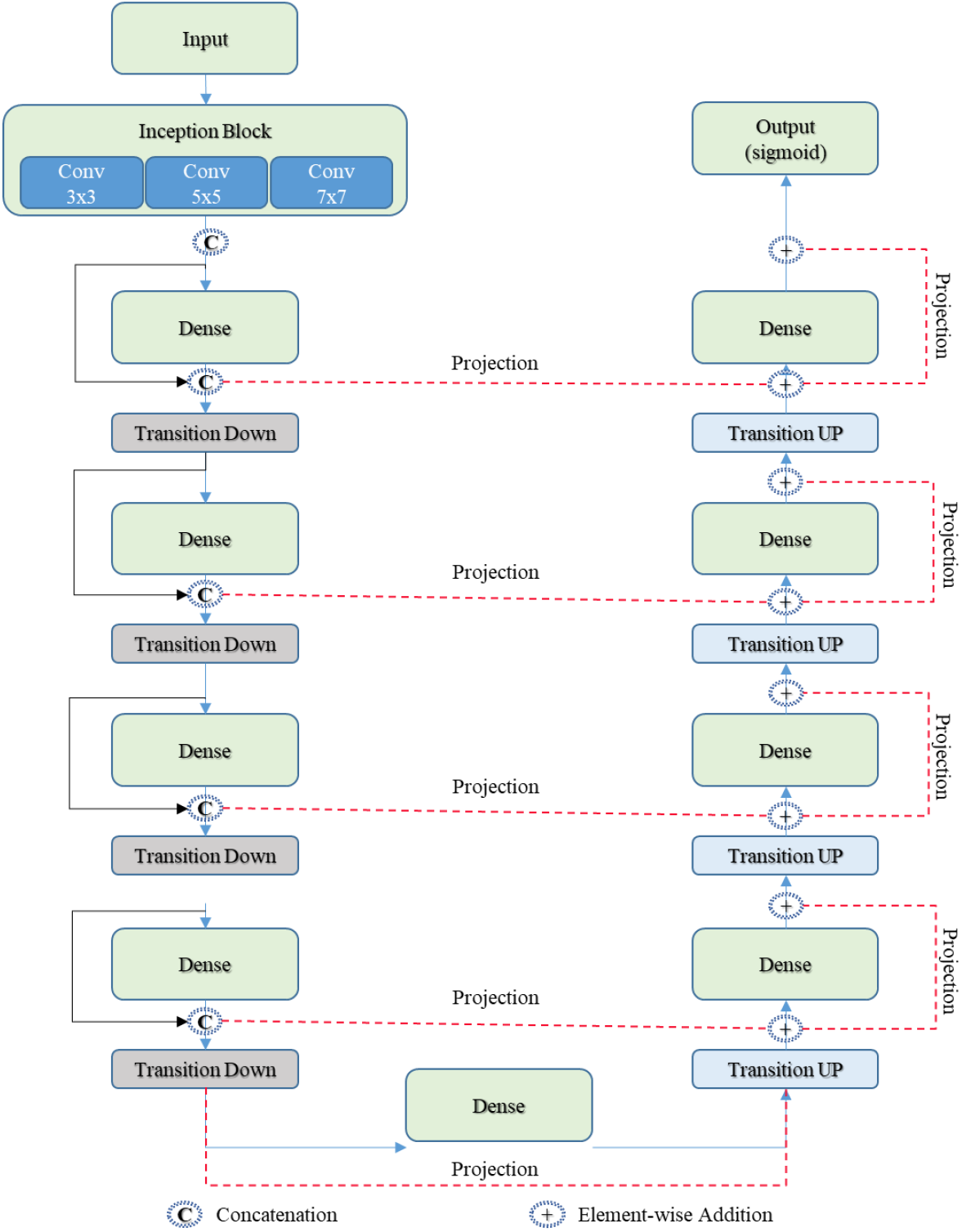
Residual Inception Densnet Architecture

**Fig. 3.**
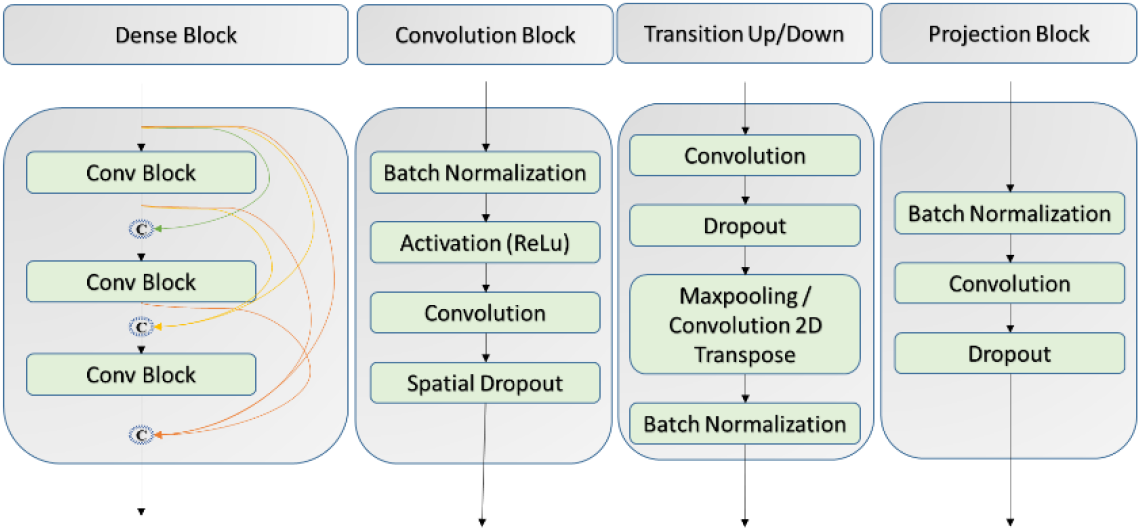
Building Blocks of Residual Inception Network. From left to right, dense block, convolution block, transition block and projection block

#### 2.2.1 Model Training and Ensemble Methodology

All of the models from the Segmentation models package were trained with full resolution axial slices of size 240×240 as input to segment the tumor subcomponents separately. The outputs for each components from the models were combined followed by post processing. The post processing step includes removing clusters of smaller size to reduce false positives. Each tumor components were then combined to form the segmentation map (Figure1.B).

The RID model was trained on 2D input patches of size 64×64. For each component of the brain tumor (e.g., hole Tumor (WT), Tumor Core (TC) and Enhancing Tumor (ET)), we trained a separate RID model with axial as well as sagittal slices as input. In addition to the six RID models, we also trained a RID with axial slices as input with patch size of 64×64 to segment TC and Edema simultaneously (TC-ED). A three dimensional RID network model was also trained to segment ET and a multiclass TC-ED (TC-ED-3D). All the models were trained with dice loss, and Adam optimizers with learning rate of 0.001 using NVIDA Tesla P40 GPU’s.

#### 2.2.2 Ensemble Methodology

The DenseNET-169, SERESNEXT-101 and SENet-154 model outputs were first combined to form segmentation maps as shown in Figure 1.B, which we will refer to as the Segmentation model output. Then, for each component we combined output from the RID models and Segmentation models output as shown in Figure.3.

**Fig. 4.**
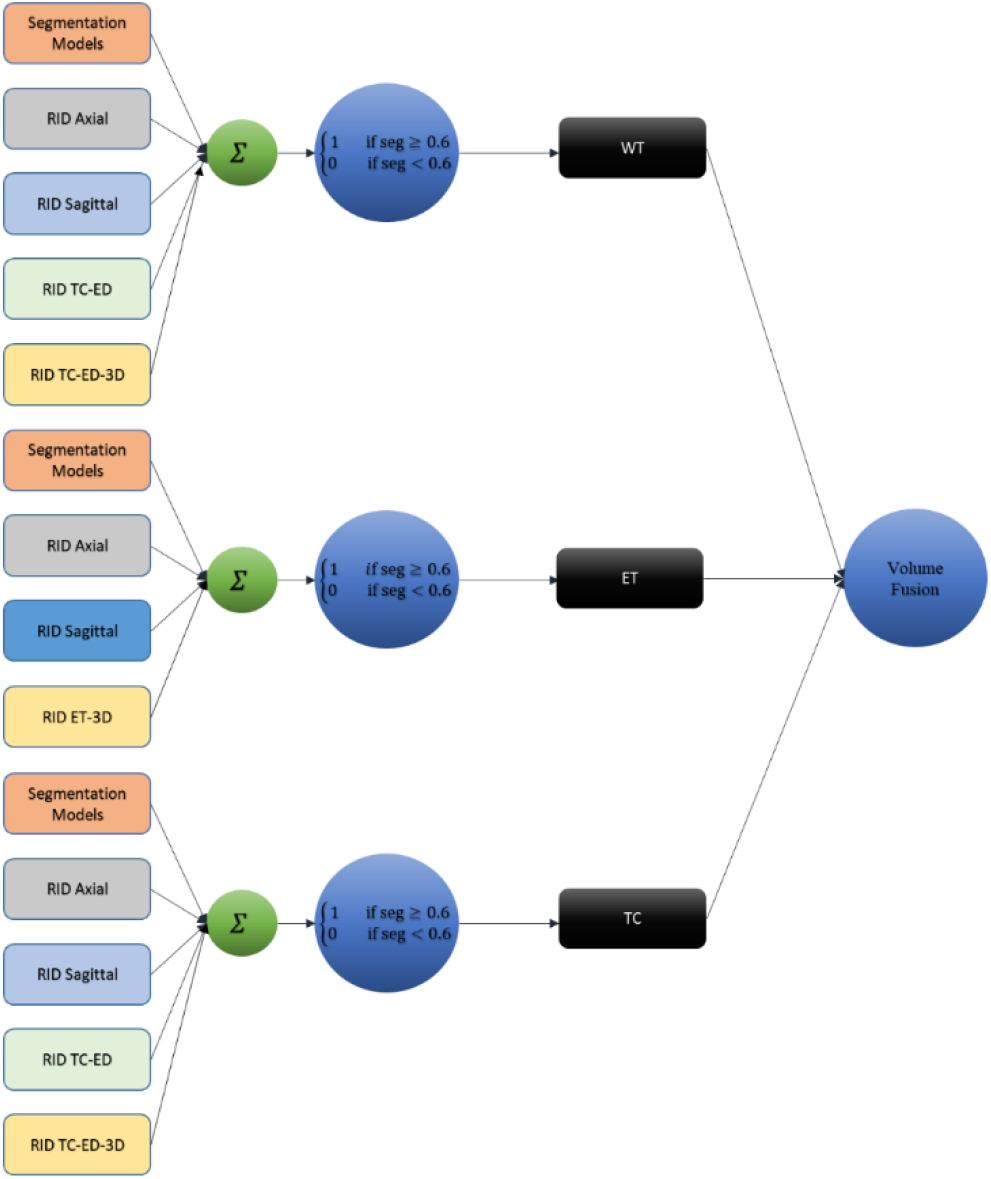
Ensemble of multidimensional and multiresolution networks. Top to bottom, ensemble for Whole Tumor (WT), Tumor Core (TC) and Enhancing Tumor (ET) respectively.

#### 2.2.3 Survival Prediction

The tumor segmentation maps extracted from the above methodology was used to extract texture and wavelet based features using the PyRadiomics (Van Griethuysen, Fedorov et al. 2017) and Pywavelets (Lee, Gommers et al. 2019) packages from each tumor subcomponent for each contrast. In addition, we added volume and surface area features of each tumor component(Feng, Tustison et al. 2018) and age. We performed feature selection based on SelectKBest features using the sklearn package(Pedregosa, Varoquaux et al. 2011, Buitinck, Louppe et al. 2013) which resulted in a reduced set of 25 features. We trained an XGBoost (XGB) model (Chen and Guestrin 2016) to classify the subjects into low (less than 300 days), medium (between 300 to 450 days) and long survivors (greater than 450 days). After predicting the class (survival range) of the subject, survival days are assigned to the subject, 150 days for the low survivor, 375 days for the mid survivor and 500 days for the high survivor category.29 subjects from the validation dataset were used to validate the trained model.

## 3 Results

### 3.1 Segmentation

The Ensemble of multiresolution 2D networks achieved 89.79%, 78.43% and 77.97% dice for WT, TC and ET respectively in validation dataset of 125 subjects.

### 3.2 Survival Prediction

Accuracy and the mean square error for the overall survival prediction for the 29 subjects using XGB was 34.5 and 122759.3 days respectively.

**Table 1.** Survival evaluation results from the xgboost (XGB) network architecture.

| Task | Network architecture | Accuracy | Mean square error |
| --- | --- | --- | --- |
| Validation | XGB | 34.5 | 122759.3 |

## 4 Discussion

We ensemble several models with multiresolution inputs to segment brain tumors. The RID network was parameter and memory efficient, and was able to converge in (as few as three epochs. This allowed us to train several models for ensemble in a short amount of time. The proposed methodology of combining multidimensional models improved performance and achieved excellent segmentation results as shown in Table.1. For survival prediction, we extracted numerous features based on texture, first order statistics and wavelets. Efficient model based feature selection allowed us to reduce the otherwise large feature set to 25 features per subject. We trained several classical machine learning models and then combined them to improve results on the validation dataset.

**Table 1.** Segmentation Results from Multiresolution 2D models and multidimension and multiresolution models

| Models | WT | TC | ET |
| --- | --- | --- | --- |
| Multiresolution 2D Models | 0.892 | 0.776 | 0.783 |
| Multidimensional and Multiresolution Models | 0.898 | 0.78 | 0.784 |

## 5 Conclusion

We demonstrate two dimensional multiresolution ensemble networks for brain tumor segmentation to generate robust segmentation of tumor subcomponents. We predicted the overall survival based on the segmented mask using an xgboost model.

